# Characteristic arrangement of nucleosomes is predictive of chromatin interactions at kilobase resolution

**DOI:** 10.1101/060327

**Authors:** Hui Zhang, Feifei Li, Yan Jia, Bingxiang Xu, Yiqun Zhang, Xiaoli Li, Zhihua Zhang

## Abstract

High-throughput chromosome conformation capture technologies, such as Hi-C, have made it possible to survey 3D genome structure. However, the ability to obtain 3D profiles at kilobase resolution at low cost remains a major challenge. Therefore, we herein report a computational method to precisely identify chromatin interaction sites at kilobase resolution from MNase-seq data, termed chromatin interaction site detector (CISD), and a CISD-based chromatin loop predictor (CISD_loop) that predicts chromatin-chromatin interaction (CCI) from low-resolution Hi-C data. The methods are built on a hypothesis that CCIs result in a characteristic nucleosome arrangement pattern flanking the interaction sites. Accordingly, we show that the predictions of CISD and CISD_loop overlap closely with chromatin interaction analysis by paired-end tag sequencing (ChIA-PET) anchors and loops, respectively. Moreover, the methods trained in one cell type can be applied to other cell types with high accuracy. The validity of the methods was further supported by chromosome conformation capture (3C) experiments at 5kb resolution. Finally, we demonstrate that only modest amounts of MNase-seq and Hi-C data are sufficient to achieve ultrahigh resolution CCI map. The predictive power of CISD/CISD_loop supports the hypothesis that CCIs induce local nucleosome rearrangement and that the pattern may serve as probes for 3D dynamics of the genome. Thus, our method will facilitate precise and systematic investigations of the interactions between distal regulatory elements on a larger scale than hitherto have been possible.

## Introduction

The 3D genome architecture underlies many cellular processes in the nucleus ^1–3^. Proximity ligation-based chromosome conformation capture (3C) and its variations ^4–6^ constitute a major engine driving the exploration of 3D genome architecture ^7,8^. Using one genome-wide version of 3C technology, Hi-C, it has been possible to explore the global 3D architecture of the human ^9,10^, mouse ^11,12^, fly ^13^ and yeast genomes ^14^. Mediator-specific 3D chromatin interaction maps have also been surveyed by the ChIA-PET method in mammals for such proteins as CTCF ^15,16^, Pol II ^16,17^, cohesin ^18^ and histone modifications ^19^. With these mappings, genomes were found to be physically separated into two compartments (A and B), one active and the other inactive ^9^. Higher resolution mapping could reveal finer structures. For example, with increased Hi-C resolution, the so-called “topologically associating domains” (TADs) ^11,20^ and sub-TAD ^21^ structures in mammals have been identified, along with detailed chromatin fiber looping structures ^16,22^. However, because most *cis*-regulatory sequences are in the size range of hundreds of base pairs and may be closely clustered ^23^, precise definition of individual enhancer-promoter interactions on a genome-wide scale remains beyond the capacity of current Hi-C methodology.

Attempts have been made to improve the resolution of chromatin-chromatin interaction (CCI) maps^15,16,22,24–29^. In order to reach one kilobase resolution, Rao et al. sequenced several billions of paired-end tags in GM12878 ^22^. Obviously, such a massive sequencing effort cannot easily be applied on a wider scale. Alternatively, capture-based methods ^15,16,24–27^, or alternative DNA cutters ^28,29^, have been described in the literature. For example, Duan and colleagues replaced restriction enzymes with DNaseI ^28^, and obtained better genome coverage and resolution than normal Hi-C. However, as a result of inherent limitations of the current 3C-based protocols, it remains challenging to substantially increase comprehensive mapping resolution to a level beyond 1∼2kb within reasonable cost constraints ^22,24^.

Chromatin 3D architecture is associated with various epigenetic features. For example, by comparing the ChIA-PET map with DNase-seq, ChIP-seq and RNAseq datasets, Snyder and colleagues showed strong association between CCIs and chromatin accessibility ^19^. Computational models have also been developed to associate histone marks with A/B compartments ^30^, CCI hubs and TADs ^31^. More recently, methods that integrate high-dimension multi-omics data in multiple cell types to predict CCIs have also been reported^32–35^. However, the massive multi-omics data required by these methods have made it difficult to elucidate the underlying mechanisms linking CCIs to chromatin dynamics.

Therefore, in this paper, we report the development of a chromatin interaction site detector (CISD) and a CISD-based chromatin loop predictor (CISD_loop) that respectively predict CCI sites and CCIs at kilobase resolution. CISD and CISD_loop only require low-resolution micrococcal nuclease digestion combined with high-throughput DNA sequencing (MNase-seq) data and low-resolution Hi-C input, respectively. We observed distinct arrangements of nucleosomes flanking the binding sites of CCI-associated factors, e.g., CTCF. Moreover, if a CCI is allele-specific, the nucleosome positioning will be more periodic in the interacting allele compared to the alternative allele. Based on these observations, it was hypothesized herein that physical CCIs could significantly alter the local chromatin context, resulting in a characteristic nucleosome arrangement pattern flanking the interaction sites. Thus, CISD was designed to predict CCI sites through the detection of such pattern, and with current annotations of TAD and low-resolution Hi-C data, CISD_loop was developed to further predict CCIs between sites predicted by CISD. We show that the predictions of CISD and CISD_loop are enriched for ChIA-PET anchors and loops, respectively. We performed 3C experiments at 5kb resolution and validated two CISD_loop predictions not reported in ChIA-PET data. Because the methods trained in one cell type can be applied to other cell types with high accuracy, the association between the characteristic nucleosome arrangement pattern and the CCI sites may be universal in human cells. The power of characteristic nucleosome arrangement pattern to predict CCIs further supports our initial hypothesis. Finally, by saturation analysis, we show that only moderate amounts of MNaseseq and Hi-C data are sufficient to achieve an ultrahigh resolution CCI map.

## Results

### Existence of distinct arrangement pattern of the nucleosomes at CCI sites

It is herein proposed that physical CCIs may alter the local chromatin context, which, in turn, causes rearrangement of the nucleosomes around the interaction sites and a resulting distinct pattern of nucleosomes. This hypothesis is based on the following facts. First, although nucleosomes have been shown to have strong DNA sequence preference ^36^, their arrangement is highly dynamic, i.e., subject to either passive remodeling by stochastically aligning to bound transcription factors (TFs) ^37^ or active remodeling by ATP-dependent remodeling enzymes ^38^. Second, genomic events or features, e.g., stably bound TF, the end of a heterochromatin domain, or simply a nucleosome-free DNA region (NFR), are sufficient to cause statistical phasing of a considerable portion of all nucleosomes ^39 37,40^. Third, the phasing patterns of the nucleosomes vary considerably among bound TFs (Fig. 1a) ^41,42^. For example, Sun and colleagues found that the nucleosome profiles of transcription factor binding sites (TFBSs) could be classified into tens of clusters, which could not be explained solely by the binding of one TF *per se*^42^. This difference may reflect the local chromatin context among TFs.

**Figure 1.**
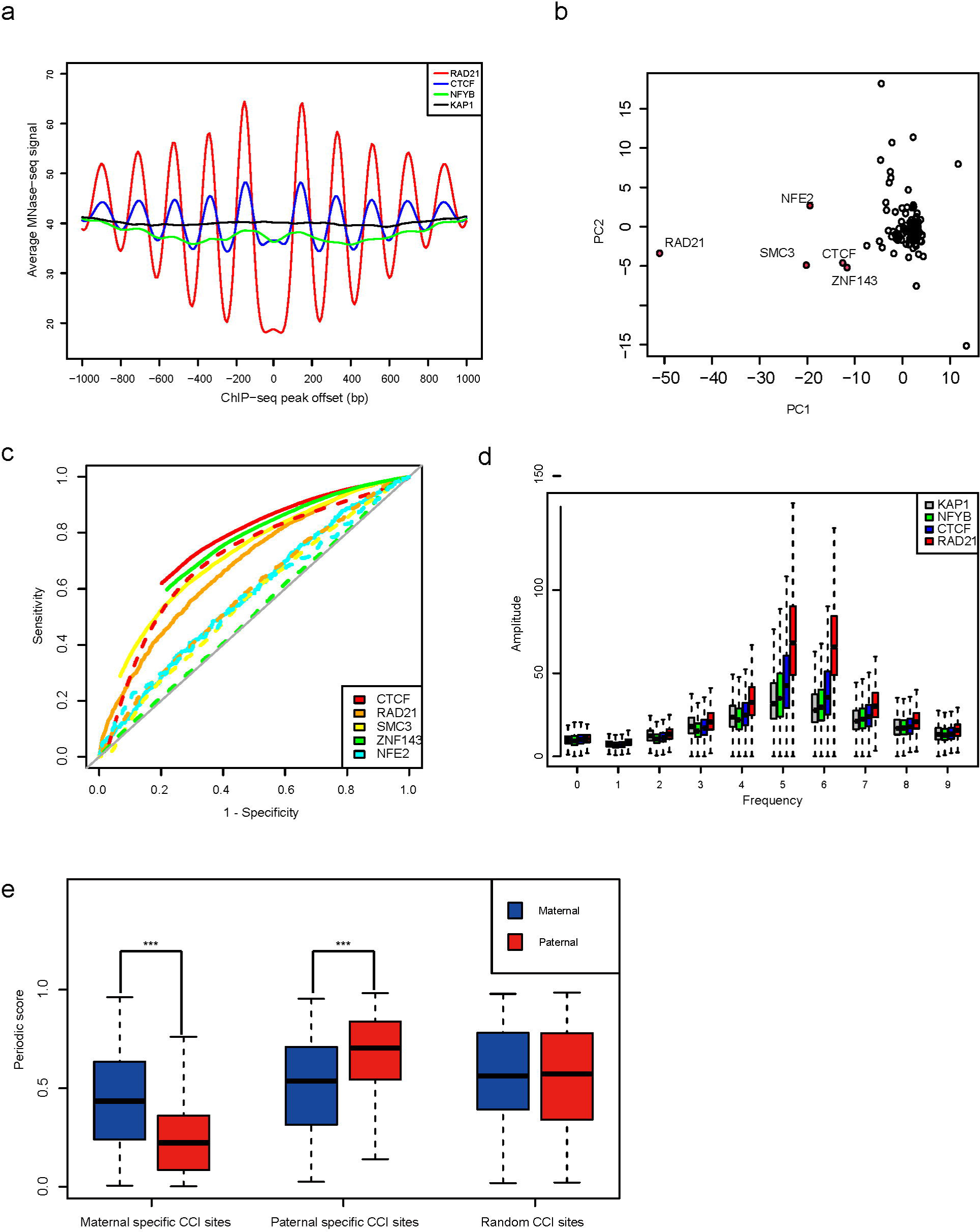
Characteristic nucleosome arrangement patterns flanking the binding sites of major CCI-mediating proteins. **(a)** The distribution of MNaseseq reads flanking the ChIP-seq peaks of Rad21, CTCF, NFYB and KAP1 in K562. **(b)** PCA analysis of MNase-seq signals flanking ChIP-seq peaks of 99 TFs. Five TFs separated from the remaining TFs are marked in red. The predictive power of ChIP-seq peaks and the periodic score of the CCI site were assessed by overlapping them with ChIA-PET loop anchors, and the ROC curves for the two methods were drawn as dashed and solid lines, respectively, in (c). **(d)** The FFT profile of MNase-seq signals flanking the ChIP-seq peaks of the four proteins in (a). (e) Aggregation analysis of allele-specific MNase-seq data at allele-specific CCI sites. Each boxplot represents the distribution of periodic scores from 200 virtual datasets. ***: rank sum test P value < 1e-10.

To test the hypothesis, we first analyzed all 99 TFs for which ChIP-seq data are available for K562 cells in the ENCODE project ^43^. Principle component analysis (PCA) of MNase-seq signals flanking the ChIP-seq peaks of the 99 TFs resulted in isolating five TFs (CTCF, RAD21, SMC3, ZNF143 and NFE2) from the others (Fig. 1b). Notably, CTCF, cohesin (RAD21 and SMC3) and ZNF143 are well known CCI-associated proteins ^18,19,44^. NFE2 is a chromatin remodeler and also reported to be engaged in enhancer-promoter interactions ^45^. However, the binding of the five TFs alone is not sufficient to predict all known CCI sites (Fig. 1c). For example, 16.3% (5,765 of 35,323) of ChIA-PET anchors do not overlap with any ChIP-seq peaks arising from these 5 TFs in K562. Therefore, to quantify the difference in nucleosome pattern arrangement (Fig. 1b), we compared the fast Fourier transform (FFT) frequency spectrum of MNase-seq data flanking the ChIP-seq peaks of 4 TFs (Fig. 1d, see Methods) and found that the amplitude of the fifth and sixth frequencies was most significantly different between CCI-associated proteins (CTCF, Rad21) and the others. Thus, by the fifth, sixth and direct component of the frequencies (see Methods), we defined a score, termed as periodic score, to index nucleosome periodicity and found the periodic score to be a better predictor of ChIA-PET anchors than the binding of TFs *per se* (Fig. 1c).

Each chromosome occupies largely mutually exclusive territories ^12,46^. Therefore, if the hypothesis is true, it can be expected that allele-specific CCI sites will also have allele-specific nucleosome arrangement patterns. Accordingly, we examined this prediction in GM12878 cells, which have well-phased, high-density SNP data^47^, and 1,707 and 1,712 maternal- and paternal-specific Rad21 ChIA-PET anchors with CTCF motifs were identified, respectively. However, because of limited SNP density flanking most of allele-specific CCI sites (+/- 1kb), allele origins could be assigned to an average of only 6.5 MNase-seq reads, making it necessary to perform an aggregation analysis (see Methods). Indeed, at the maternal-specific CCI sites, we found that nucleosomes are more periodically arranged in the maternal allele compared to the paternal allele. Meanwhile, the pattern is just the opposite at the paternal-specific CCI sites, and no significant difference can be found in randomly selected CCI sites (Fig. 1e). Taken together, this evidence supports our hypothesis that nucleosome arrangement pattern is reflective of the local chromatin environment and might be utilized for the detection of CCIs.

### Detecting chromatin interaction sites at kilobase resolution with CISD

Therefore, based on the predictive potential of nucleosome arrangement in the context of local chromatin environment, as shown above, we developed the chromatin interaction site detector (CISD) to identify CCI sites. For any given genome locus, CISD takes MNase-seq data as input and determines whether the nucleosomes display the assumed arrangement pattern of a CCI site. First, CISD is composed of a logistic regression model (LRM) that determines whether the input genome locus has a periodic nucleosome arrangement pattern. If so, a second support vector machine (SVM) model then determines whether the locus has a nucleosome pattern characteristic of CCIs (hereinafter termed CISD sites when predicted by CISD) (Fig. 2a). The AUC for the LRM was 0.97 and 0.92 in K562 and GM12878 cells, respectively, and five-fold cross-validations of the accuracy of the SVM model were above 80% in both cell types. The resolution of CISD is defined as the length of the genome segment needed to make a credible prediction. In this work, we took 1kb as the default resolution. Using the default threshold for the periodic score (0.5), we applied CISD to K562 and GM12878 cells and predicted 22,112 and 26,801 CISD sites, respectively. Some canonical CCI sites have been successfully revealed by CISD, such as human β-globin locus in K562 (Supplementary Fig. 1a). The genome-wide distributions of the CISD sites are similar between the two cell types (Supplementary Fig. 1b).

**Figure 2.**
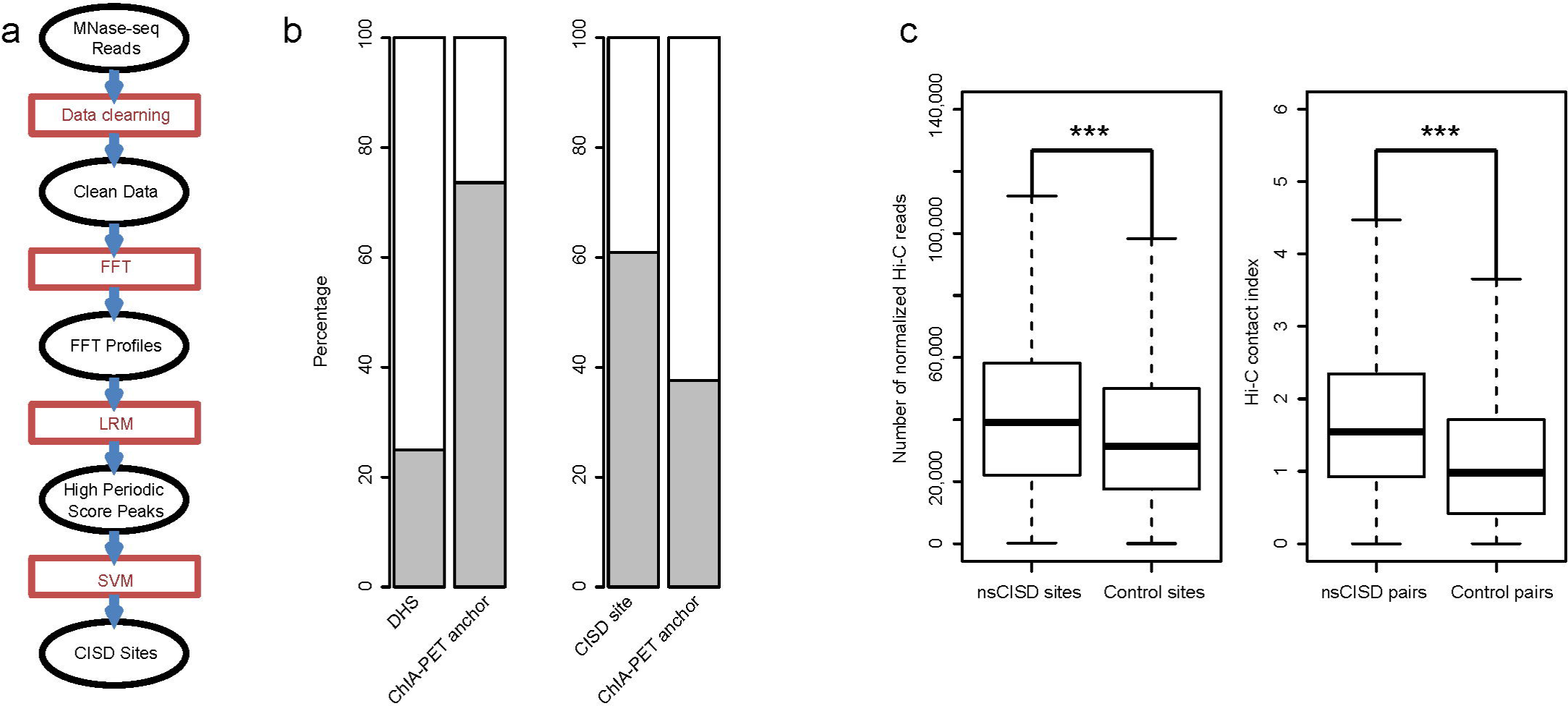
CISD workflow and performance. **(a)** The flow chart of CISD. The ovals represent datasets that were used or generated, and the square boxes represent data processing steps. **(b)** Fractions of DHSs and CISD sites that overlap ChIA-PET loop anchors (left columns) in K562 and fractions of ChIA-PET loop anchors that overlap DHSs and CISD sites (right columns). **(c)** Distribution of Hi-C reads count around nsCISD and control sites (left); distribution of Hi-C contact index between nsCISD sites and between control sites (right) in K562. For each site, the reads count is calculated from the normalized Hi-C contact matrix. The 5kb resolution matrix was used in this figure. (***: rank sum test P-value< 2.2e-16).

To further evaluate CISD, we took ChIA-PET loop anchors as the gold standard because ChIA-PET is believed to identify CCIs at high resolution^48^. Compared to DNase I hypersensitive sites (DHSs), which were reported to be predictive of chromatin looping anchors ^19,35,49^, CISD sites enriched more ChIA-PET anchors. In K562, CISD sites were 2.4-fold more enriched for ChIA-PET anchors than were DHSs, as 24.9% (43,340 out of 174,043) and 60.0% (13,278 out of 22,112) of DHSs and CISD sites, respectively, overlapped the ChIA-PET loop anchors (Fig. 2b). Results largely corresponding to these were also seen in GM12878 cells (Supplementary Fig. 1c). In the LRM model, when the threshold for periodic score was increased from 0.5 to 0.9, only 12.6% more CISD sites (up from 60.0% to 72.6%) overlapped ChIA-PET loop anchors in K562, and the corresponding figure for GM12878 was only 21.1% (up from 72.3% to 93.4%), suggesting that the predictive power of CISD is not sensitive to the threshold for the periodic score it used. Because the DHSs are ubiquitous in the genome, it is not surprising to find that the total number of DHSs overlapping ChIA-PET anchors is larger than the total number of CISD sites (Fig. 2b, Supplementary Fig. 1c).

Based on this disparity, we decided to examine to extent to which CISD sites lacking support from Chl-PET data, termed as nonsupported (ns) CISD sites, might be involved in chromatin interactions. To accomplish this, we compared the number of Hi-C reads around the nsCISD sites to those around genomic regions that have a high periodic score, but were not predicted as CISD sites. Indeed, the nsCISD sites were not only significantly more enriched for Hi-C reads than were control sites (P-value<2.2e-16, rank sum test), but a considerably higher enrichment of Hi-C-mated reads was observed between nsCISD sites than that observed between control sites (P-value <2.2e-16, rank sum test, Fig. 2c, Supplementary Fig. 1d). Finally, we experimentally validated two nsCISD sites by 3C at 5kb resolution, as detailed below.

### Prediction of CCIs between CISD sites with CISD_loop

To predict chromatin loops between CISD sites, we developed CISD_loop which takes CISD sites and low-resolution Hi-C data as input. Briefly, CISD_loop is a SVM model trained on intra-TAD CISD site pairs taking the Hi-C contact index and the distance between the two anchors as features (see Methods and Fig. 3a,b). The 5-fold cross-validation of accuracy of this SVM predictor was 79.0% and 76.6% in K562 and GM12878 cells, respectively. We applied CISD_loop on Hi-C data from K562 and GM12878 cells and predicted 35,143 and 60,342 interactions, respectively, out of which 14.4% and 21.3% could be supported by ChIA-PET loops. Compared to random intra-TAD CISD site pairs, CISD_loop predictions have 3- and 2.3-fold higher enrichment for ChIA-PET loops in K562 and GM12878 cells, respectively.

**Figure 3.**
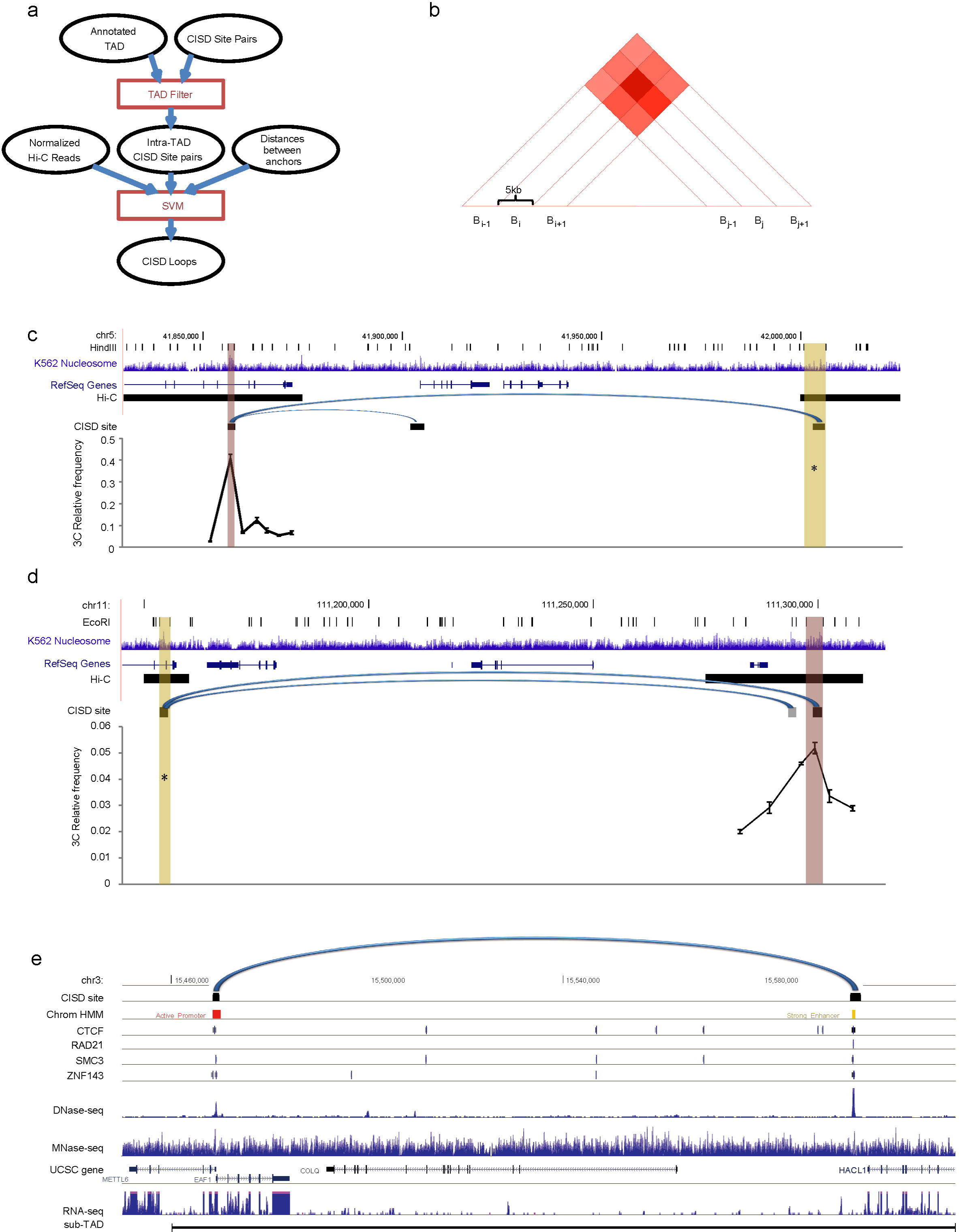
CISD_loop workflow and performance. **(a)** Flowchart of CISD_loop. For explanation, see Figure 2a. **(b)** Illustration of the Hi-C contact index calculation (see Methods). The diamonds represent the entries in the normalized Hi-C contact matrix. Two 3C experiments were performed **(c)** between chr5:42,000,785-42,006,207 (anchor) and chr5:41,856,243-41,857,738 (target) and **(d)** between Chr11:111,153,407-111,155,719 (anchor) and Chr11:111,297,381-111,301,207 (target). Each data point represents mean ± SEM of three technical replicates and three normalized biological replicates. The CCIs predicted by CISD_loop are marked as arches. The third CISD site was marked in grey, indicating that it could not be validated in this experiment because a restriction site appears in the site. **(e)** A putative chromatin interaction predicted by CISD_loop is marked as an arch. MNase-seq reads, ChIP-seq peaks, DNase-seq reads and CISD_loop predictions are shown within a sub-TAD region on chr3:15,460,000-156,200,00.

To assess the reliability of CISD_loop predictions that have no ChIA-PET data support, we performed 3C experiments to validate two such examples (Fig. 3c,d). The two were selected because the restriction sites flanking the CISD sites had suitable density to allow accurate determination of the interaction sites within an approximate 5kb window. The restriction fragments with nsCISD sites had the highest crosslinking frequencies, and all qPCR products were confirmed to be the expected ligation products using Sanger sequencing. In comparison to more than 10kb length of the Hi-C segments in which the two selected “nsCISD” sites were located, 3C experiments confirmed our predictions at 5kb resolution (Fig. 3c,d).

Evidence for the reliability of CISD_loop predictions also comes from transcriptome data. For example, in a sub-TAD (chr3:15,460,000-156,200,00; Fig. 3e)^22^, a strong enhancer and three genes (METTL6, EAF1 and COLQ) are annotated in the UCSC genome browser. However, RNA-seq data show that only METTL6 and EAF1 are actively transcribed in K562 cells. The promoter of the transcriptionally silent gene COLQ is much closer to the enhancer than the common bidirectional promoter of METTL6 and EAF1. This apparent anomaly could be explained by our CISD_loop prediction which showed direct contact between the strong enhancer and bidirectional promoter of METTL6 and EAF1, while bypassing COLQ (Fig. 3e).

### The characteristic nucleosome pattern flanking CCI sites is concordant in cell types

To test whether the characteristic nucleosome pattern is concordant across cell types, we trained CISD and CISD_loop with data from one cell type and tested them on data from another cell type (Table 1). When the training data were from K562 cells, both CISD and CISD_loop had comparable predictive power in K562 and GM12878 cells, and the CISD_loop predictions in GM12878 had an even higher validation rate by the ChIA-PET data than that of K562 cells. When the training data were from GM12878, the intercell-type validation rate was substantially reduced. This could be explained by comparing the ChlA-PET GM12878 and K562 datasets. The ChIA-PET GM12878 contains more Pol II-mediated CCIs compared to that of K562, which introduces training biases in the CISD/CISD_loop (73.6% and 17.9% ChIA-PET data were targeting Pol II, in GM12878 and K562, respectively). Even so, on average, validation rates of 46.3% and 6.2% were still 1.9-fold and 4.4-fold higher than random assignments for CISD and CISD_loop, respectively, compared to 25.0% and 1.4%. Thus, the underlying nucleosome pattern that CISD identified with data from K562 cells may be more representative of a possible “consensus nucleosome pattern” in cells. In accordance with this surmise, the default CISD and CISD_loop were therefore trained with K562 data.

**Table 1.** Intra- and intercell-type performance of CISD and CISD_loop.

| Training set | Testing set | # of CISC sites | % of ChIA-PET anchors | # of CISC loops | % of ChIA-PET loops |
| --- | --- | --- | --- | --- | --- |
| K562 | K562 | 22,112 | 62.30% | 35,143 | 14.40% |
| K562 | GM12878 | 17,441 | 59.90% | 16,681 | 39.70% |
| GM12878 | GM12878 | 26,801 | 72.30% | 60,342 | 21.30% |
| GM12878 | K562 | 34,036 | 46.30% | 117,608 | 6.20% |

### Modest amounts of data are sufficient for CISD/CISD_loop to achieve ultrahigh resolution predictions

To examine how many MNase-seq reads are necessary to obtain highly accurate CISD site predictions, we composed testing datasets by randomly sampling descending numbers of mapped MNase-seq reads, e.g., one half, one quarter, and one eighth, from the original data in chromosome one of K562 cells, or about 142 million mapped reads ^41^. The original sequencing depth was about 20-fold, and the testing datasets simulated sequencing depths of 10-, 5- and 2.5-fold, with read density equivalent to about 570, 285, 143, and 71 kilo reads per million base-pair region, respectively. Even with the lowest number of reads, CISD could successfully identify periodic nucleosome regions (Fig. 4a), and over one third of the predictions overlapped with currently available ChIA-PET loop anchors (Fig. 4b). However, because the overlapping proportion dropped substantially when read coverage became less than 5-fold, we suggest 5-fold or higher coverage for CISD site prediction.

**Figure 4.**
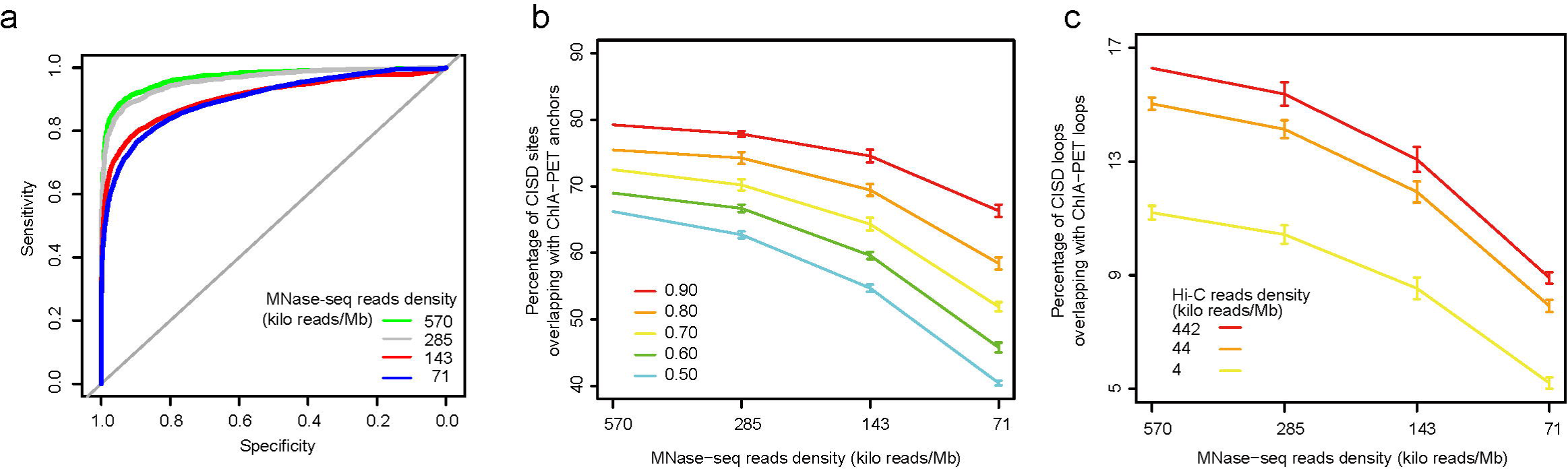
MNase-seq and Hi-C data requirements for high-resolution prediction by CISD and CISD_loop. **(a)** ROC curves for LRM predictions with different portions of MNase-seq data. **(b)** Percentages of CISD sites supported by ChIA-PET loop anchors under different periodic score thresholds and different portions of MNase-seq data. **(c)** Percentages of CISD_loop predictions supported by ChIA-PET loops with different portions of MNase-seq and Hi-C data. The threshold of periodic score in the LRM step was set at 0.5. Each data point represents mean ± SD of 10 technical replicates.

We next confirmed how many MNase-seq and Hi-C reads would be needed to obtain high accuracy of CISD loops. We composed the testing data with all, 10% and 1% of the current Hi-C reads in chromosome one of K562 cells^22^, with read density equivalent to about 442, 44 and 4 kilo reads pre million base-pair region, respectively, and examined the performance of CISD_loop for all combinations of the three Hi-C and four MNase-seq testing datasets (Fig. 4c). Exponential reduction of Hi-C reads number did not substantially affect the validation rate. With only 10% of Hi-C reads, CISD_loop still had nearly the same validation rate as it did with the full data, and the validation rate only dropped to half when the Hi-C data were reduced to 1%. Thus, we may set the sequencing depth according to the desired validation rate. For example, if 10% of CISD loops are expected to be supported by ChIA-PET, then the combination of 20-fold coverage of MNase-seq and about 1% of Hi-C reads (equivalent to about 570 and 4 kilo reads/Mb for MNase-seq and Hi-C, respectively) or, alternatively, 5-fold coverage of MNase-seq and 10% Hi-C reads (equivalent to about 143 and 44 kilo reads/Mb for MNase-seq and Hi-C, respectively), would allow CISD/CISD_loop to achieve the desired validation rate. Finally, we also trained the model in one cell type, either K562 or GM12978, and tested it in the other cell type and obtained similar results (Supplementary Fig. 2a-c), suggesting that the method can be widely applied in human cell lines.

## Discussion

In this paper, we developed CISD/CISD_loop for genome-wide identification of potential CCI sites and loops at kilobase resolution. This ultrahigh resolution can be achieved because the number of nucleosomes with distinct arrangement pattern flanking barriers is not large ^50^, and current MNase-seq data are sufficient to detect such pattern. In addition to ultrahigh resolution, our methods also make 3D genome profile exploration more economical than ultra-deep sequencing, as only MNase-seq and low-resolution Hi-C data are used.

As a complement to current methods, e.g., EpiTensor ^33^ and TargetFinder ^34^, which detect consistent CCIs across cell types, CISD/CISD_loop can predict CCIs in a cell type-specific manner, essentially because our methods do not rely on data from other cell types. Cell type-specific CCI prediction is an important advance, given the highly dynamic nature of chromatin interactions and prevalence of cell-type specificity on promoter-enhancer interactions ^3,37,38^.

Epigenetic features of the genome have been used for computational modeling of chromatin architecture ^19,30,31,33,34^. The success of such models suggests the presence of profound links between CCI sites and the dynamics of the chromatin *per se* ^34^. However, data integration-based methods can only provide limited insight toward the elucidation of such links. The predictive power of CISD/CISD_loop suggests that CCIs may serve as special barriers which alter the local chromatin context and cause rearrangement of the nucleosomes. This, in fact, may be a potential mechanism linking CCI sites and the dynamic behavior of nucleosomes. Although the actual characteristic pattern is complicated and cannot be simplified as nucleosomal periodicity or openness of chromatin, the pattern per se is universal cross cell types.

The biochemical mechanisms underlying chromatin interactions are far from being understood. It is certain to be a complex process with many factors involved at multiple levels. Thus, any method relying on a single datum is likely to yield an incomplete result, and indeed, the sensitivity of CISD is limited. Therefore, it is possible that further improvement may be achieved by marrying data integration and hypothesis-driven modeling. Furthermore, as indicated by Whalen et al., the information relevant to looping interactions is not just limited to the interacting loci ^34^. Thus, taking the data from outside the interacting loci into account also merits further investigation.

The considerable attention in recent literature directed towards 3D genome studies reflects the importance of such knowledge. CISD and CISD_loop provide an approach that facilitates the expansion of the field of 3D genome research by allowing the exploration of more cell types, tissues, and species.

## Method and Materials

### Cell culture and 3C experiment

Human K562 cells were obtained from ATCC (http://www.atcc.org/) and were grown in RPMI 1640 medium containing 10% FBS and penicillin/streptomycin. Cells were routinely tested for mycoplasma contamination and were characterized by DNA fingerprinting analysis. The 3C assay was performed as previously described with a few modifications ^9,22^. Briefly, 5 million cells were crosslinked with 1% formaldehyde for 10min. After cells were lysed, DNA was digested with HindIII or EcoRI overnight with rotation. In-nucleus ligation was conducted using T4 ligase in order to reduce spurious contact from random ligation. Crosslinking was reversed, and DNA was purified by two rounds of phenol-chloroform extraction and ethanol precipitation. DNA concentrations were measured using a Qubit fluorometer and subsequently diluted for real-time quantitative PCR with SYBR green. BAC clones spanning the analyzed loci (RP11-48E24 and 241N21 for EcoRI loci; RP11-100621 and 997N5 for HindIII loci) were digested, religated and used as a control for primer efficiency. To correct for the differences in quality and quantity of templates in different replicates, the ligation frequencies were normalized to control interactions at the housekeeping Ercc3 gene locus. A full list of primers is available upon request.

### Data

We downloaded the MNase-seq ^41^, ChIP-seq^51^, DNase I hypersensitive sites ^52^, and ChIA-PET data (GSE39495) in the ENCODE project^43^. GEO accession numbers include GSM920557; GSM920558; GSM736629; GSM736566; GSM736496; GSM736620; GSM970216; GSM935310; GSM935311; GSM935319; GSM935336; GSM935337; GSM935338; GSM935340; GSM935343; GSM935344; GSM935355; GSM935356; GSM935358; GSM935361; GSM935368; GSM935371; GSM935372; GSM935373; GSM935374; GSM935385; GSM935388; GSM935391; GSM935392; GSM935394; GSM935401; GSM935402; GSM935407; GSM935410; GSM935411; GSM935414; GSM935425; GSM935428; GSM935429; GSM935433; GSM935439; GSM935464; GSM935466; GSM935467; GSM935468; GSM935469; GSM935470; GSM935471; GSM935472; GSM935473; GSM935474; GSM935475; GSM935479; GSM935481; GSM935487; GSM935488; GSM935490; GSM935494; GSM935495; GSM935496; GSM935497; GSM935499; GSM935501; GSM935502; GSM935503; GSM935504; GSM935505; GSM935516; GSM935520; GSM935521; GSM935532; GSM935539; GSM935540; GSM935544; GSM935546; GSM935547; GSM935548; GSM935549; GSM935565; GSM935568; GSM935569; GSM935573; GSM935574; GSM935575; GSM935576; GSM935594; GSM935595; GSM935597; GSM935598; GSM935599; GSM935600; GSM935602; GSM935616; GSM935632; GSM935633; GSM935634; GSM935642; GSM935645; GSM1003608; GSM1003609; GSM1003610; GSM1003611; GSM1003620; GSM1003621; GSM1003622; GSM1003625. Additional ChIA-PET data were downloaded from the studies of ^19^ and ^16^. Hi-C data were downloaded from ^11,22^.

### Determination of allele-specific MNase-seq reads

We downloaded the most updated phased SNP of the GM12878 cell line in the 1000 Genomes Project ^47^, from Gerstein’s lab ^53^. By overlapping the mapped reads with phased SNPs, we identified 8,382,815 and 8,456,093 paternal-specific and maternal-specific MNase-seq reads, respectively.

### Determination of allele-specific CCI

We took the ChIA-PET loop anchors in the GM12878 cell line as the gold standard CCI sites^48^. However, the anchor lengths were too short (mean = 424bp) to carry out a sufficient number of SNPs. Therefore, instead of using ChIA-PET data, we pooled all Hi-C reads that mapped within a 5kb genome region flanking each ChIA-PET loop anchor ^22^. We used the ratio of the maternal-specific read number to the paternal-specific read number to index the allele specificity of the CCI in each region. We did not consider CCIs in chromosome X. As the index has a bell shape distribution (Supplementary Fig. 3), we took the first and last 10% as maternal- and paternal-specific CCIs anchors, respectively. We further filtered CCIs which had an abnormally high number of reads.

### Aggregation analysis on allele-specific MNase-seq data

The purpose of this analysis is to aggregate information from sporadic allele-specific MNase-seq reads to reveal a general pattern of nucleosome arrangement at allele-specific CCI sites. As an example, we describe how the information of nucleosome arrangement in the maternal allele at paternal-specific CCI sites is aggregated. We first collected maternal-specific reads in the 10kb flanking region of paternal-specific CCI sites, denoted as *ASR*(*m,p*). By sampling 5500 reads with replacement from the *ASR*(*m,p*), we could generate a virtual allele-specific sequencing dataset. We aggregated reads in this dataset centered by the CTCF motifs found in each CCI site. The periodic score could then be calculated as described below. We generated 200 virtual datasets for *ASR*(*m,p*) and drew boxplot of periodic scores as in Figure 1e.

### Chromatin interaction site detector (CISD)

Basically, CISD determines whether the nucleosome arrangement pattern in a given genome locus is a pattern that is characteristic of chromatin interactions. The algorithm can be largely separated into a data preparation section and two model training sections (Fig. 2a).

#### Data preparation

Here, we convert MNase-seq data into the frequency spectrum by fast Fourier transform (FFT). CISD first smoothes the input MNaseseq reads. For any given genomic region (1kb-long in this study), the mapped reads are binned into 10bp-long bins, resulting an *n-* dimensional vector *V*. *V* is then fed into the iNPS for denoising and smoothing ^54^. The iNPS is an improved version of NPS ^55^; both iNPS and NPS denoise and smooth the wave-form signal by Laplacian of Gaussian convolution (LoG) ^54,55^. After the LoG, the frequency feature of the original data is preserved, while its direct current component is substantially reduced. The denoised and smoothed *V* (denoted as *V’*) is further normalized by dividing the standard deviation of *V’* over the whole genome, and it was denoted as 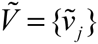, *j* = 0,2,…,n − 1. After data denoising, smoothing and normalization, CISD converts the linear data into frequency space. To do this, for any given 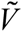, FFT is applied to retrieve the frequency information. In general, FFT is a fast computational method for Discrete Fourier Transformation (DTF). The DTF converts an *n*-dimensional vector 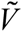 of complex numbers into a complex number vector *C* = {*c*_*j*_}, *j* = 0,2,3,…, *n* − 1,

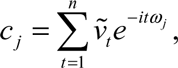

where *i* is the basic unit of the imaginary number, and *ω*_*j*_ = 2*πj/n*. Since 1) *C* is conjugate symmetric and 2) we are only interested in the modulus of *C*, we discarded that half of the elements in *C* where the real parts are negative. We thus arrive at the definition of the FFT profile (denoted as *F*) of the nucleosome arrangement in a genome segment such that

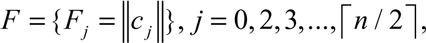

where ‖*c*_*j*_‖ denotes the modulus of *c*_*j*_.

#### Model training one: Periodic region detection

We define a metric for the periodicity of a given genomic region by a logistic regression model (LRM). To train the LRM, we constructed a positive and a negative dataset containing CTCF/cohesin co-binding of the ChIA-PET anchors and the control genomic regions, respectively, in the GM12878 and K562 cell types. In the positive dataset, the cobinding of CTCF and RAD21 is inferred by the ChIA-PET anchors. CTCF and cohesin are considered to be co-binding if the two ChIA-PET anchors overlap by more than one basepair. The negative set is composed of genomic segments that are located at least 5kb away from the CTCF, cohesin and ZNF143 binding sites and are also not in promoter regions, i.e., at least 5kb away from UCSC annotated transcription start sites. We chose *F*_0_, *F*_5_ and *F*_6_ in the FFT profiles (*F*_*j*_) as the features with which to train the LRM. The LRM was trained by R. An artificial threshold of LRM score (**periodic score**) was then chosen to determine if the input genomic segment is carrying a periodic nucleosome pattern. In this paper, we took 0.5 as the threshold and applied the LRM to the whole genome, denoting the determined periodic nucleosomal regions as high score peaks (**HSPeaks**) to be used as input for the next step.

#### Model training two: interaction site detection

As we have shown above, not all periodic nucleosomal regions are associated with chromatin interactions. Accordingly, we trained a supporting vector machine (SVM) to further distinguish interactive loci from the remaining periodic nucleosomal regions. We took the full frequency spectrum (*F*) as the feature. To train the SVM, we constructed a positive and a negative dataset from the HSPeaks. The positive sets consisted of overlapping CTCF and cohesin ChIA-PET anchors, while the negative set was randomly sampled from a subset of HSPeaks that did not overlap any ChIA-PET anchors. The SVM model was implemented by R-package “e1071” with default parameter settings.

### CISD-based Chromatin loop predictor (CISD_loop)

CISD_loop is a SVM model for determination of intra-TAD chromatin loops between CISD sites. In addition to CISD, TAD annotation and raw Hi-C reads are required for the operation of CISD_loop. The TADs annotated in hESC were used as the reference for all human cells, as the TAD structure is believed to be largely consistent among tissue types ^11^. CISD_loop was trained according to the following procedure. First, we constructed a “total” dataset composed of all intra-TAD CISD site pairs. Then, the training and testing datasets were drawn from the “total” dataset so that the positive set (5,000 data points in this work) was composed of the CISD pairs that overlapped with ChIA-PET loops, and the negative set was an identical number of randomly sampled CISD pairs from the remaining data. Two features were employed in CISD_loop, namely, the Hi-C contact index and the distances between the pairs of CISD sites (Fig. 3b).The Hi-C contact index was defined using the normalized Hi-C contact matrix with a genome bin size of 5kb ^22^. For any given pair of CISD sites, (*C_i_*, *C_j_*), we denoted the genome bins that cover the CISD sites *C_i_* and *C_j_* as *B_i_* and *B_j_*, respectively. We also denote *B_i-1_* and *B_i+1_* as the 3’- and 5’-neighbor bins of *B_i_*, respectively. Then, the Hi-C contact index was defined as the average number of the 3×3 square [*B_i-1_* − *B_i+1_* × *B_j-1_* − *B_j+1_*] in the Hi-C contact matrix (Fig. 3b). The SVM model was implemented by R-package “e1071” with default parameter settings.

#### Code availability

Source code for the CISD and CISD_loop can be found at https://github.com/huizhangucas/CISD.

#### Evaluation of the models

In all total datasets, we had identical numbers of entries in the positive and negative sets. The accuracy of a model was defined as (TP + TN) / (TP + TN + FP + FN), where TF, TN, FF, and FN represent true positives, true negatives, false positives and false negatives, respectively. Standard five-fold cross-validation was performed according to the following procedure. The original total data sample was randomly partitioned into 5 subsets of equal size. Of the five subsets, a single subset was retained as the testing data, and the remaining 4 subsets were used as training data. We repeated this procedure 5 times with each of the 5 subsets used exactly once as the testing data. The 5 results were averaged for the final evaluation.

## Author Contributions

ZZ conceived this project. HZ, ZZ developed CISD and CISD_loop. FL designed the 3C experiment. FL, YJ and XL performed the 3C experiment. HZ, BX, YZ analyzed data, ZZ, HZ and FL prepared the manuscript. All authors read and approved the final manuscript.

## Acknowledgments

We thank Dr. Zhihu Zhao for the help in setting up the 3C experiment. We thank Dr. Qianfei Wang and Dr. Changqing Zeng for the help in accessing experimental equipment, and we appreciate helpful discussions with Dr.Chengqi Wang, Lijia Yu and Guangyu Wang. We thank Dr. Geir Skogerbo and Mr. David Martin for the language correction on the manuscript. We also acknowledge the ENCODE Consortium and the ENCODE production laboratories that generated the datasets provided by the ENCODE Data Coordination Center used in this manuscript. This work was supported by grants from the National Nature Science Foundation of China (NSFC, 91540114, 31271398), the National High Technology Development 863 Program of China (2014AA021103), and Special Program for Applied Research on Super Computation of the NSFC-Guangdong Joint Fund (the second phase) to ZZ. The funders had no role in study design, data collection and analysis, decision to publish, or preparation of the manuscript.

## References

1. Roy, A.L., Sen, R. & Roeder, R.G. Enhancer-promoter communication and transcriptional regulation of Igh. Trends Immunol 32, 532–539 (2011).

2. Li, G. et al. Extensive promoter-centered chromatin interactions provide a topological basis for transcription regulation. Cell 148, 84–98 (2012).

3. Zhang, Y. et al. Chromatin connectivity maps reveal dynamic promoter-enhancer long-range associations. Nature 504, 306–10 (2013).

4. de Wit, E. & de Laat, W. A decade of 3C technologies: insights into nuclear organization. Genes Dev 26, 11–24 (2012).

5. de Laat, W. & Dekker, J. 3C-based technologies to study the shape of the genome. Methods 58, 189–91 (2012).

6. Simonis, M., Kooren, J. & de Laat, W. An evaluation of 3C-based methods to capture DNA interactions. Nat Methods 4, 895–901 (2007).

7. Belton, J.M. et al. Hi-C: a comprehensive technique to capture the conformation of genomes. Methods 58, 268–76 (2012).

8. Zhang, Y. et al. Spatial organization of the mouse genome and its role in recurrent chromosomal translocations. Cell 148, 908–21 (2012).

9. Lieberman-Aiden, E. et al. Comprehensive mapping of long-range interactions reveals folding principles of the human genome. Science 326, 289–93 (2009).

10. Naumova, N. et al. Organization of the mitotic chromosome. Science 342, 948–53 (2013).

11. Dixon, J.R. et al. Topological domains in mammalian genomes identified by analysis of chromatin interactions. Nature 485, 376–80 (2012).

12. Selvaraj, S., J, R.D., Bansal, V. & Ren, B. Whole-genome haplotype reconstruction using proximity-ligation and shotgun sequencing. Nat Biotechnol 31, 1111–8 (2013).

13. Sexton, T. et al. Three-dimensional folding and functional organization principles of the Drosophila genome. Cell 148, 458–72 (2012).

14. Duan, Z. et al. A three-dimensional model of the yeast genome. Nature 465, 363–7 (2010).

15. Handoko, L. et al. CTCF-mediated functional chromatin interactome in pluripotent cells. Nat Genet 43, 630–8 (2011).

16. Tang, Z. et al. CTCF-Mediated Human 3D Genome Architecture Reveals Chromatin Topology for Transcription. Cell 163, 1611–27 (2015).

17. Sandhu, K.S. et al. Large-scale functional organization of long-range chromatin interaction networks. Cell Rep 2, 1207–19 (2012).

18. Demare, L.E. et al. The genomic landscape of cohesin-associated chromatin interactions. Genome Res 23, 1224–34 (2013).

19. Heidari, N. et al. Genome-wide map of regulatory interactions in the human genome. Genome Res 24, 1905–17 (2014).

20. Nora, E.P. et al. Spatial partitioning of the regulatory landscape of the Xinactivation centre. Nature 485, 381–5 (2012).

21. Phillips-Cremins, J.E. et al. Architectural Protein Subclasses Shape 3D Organization of Genomes during Lineage Commitment. Cell 153, 1281–95 (2013).

22. Rao, S.S. et al. A 3D Map of the Human Genome at Kilobase Resolution Reveals Principles of Chromatin Looping. Cell 159, 1665–80 (2014).

23. Shen, Y. et al. A map of the cis-regulatory sequences in the mouse genome. Nature 488, 116–20 (2012).

24. Naumova, N., Smith, E.M., Zhan, Y. & Dekker, J. Analysis of long-range chromatin interactions using Chromosome Conformation Capture. Methods 58, 192–203 (2012).

25. Kolovos, P. et al. Targeted Chromatin Capture (T2C): a novel high resolution high throughput method to detect genomic interactions and regulatory elements. Epigenetics Chromatin 7, 10 (2014).

26. Dryden, N.H. et al. Unbiased analysis of potential targets of breast cancer susceptibility loci by Capture Hi-C. Genome Res 24, 1854–68 (2014).

27. Hughes, J.R. et al. Analysis of hundreds of cis-regulatory landscapes at high resolution in a single, high-throughput experiment. Nat Genet 46, 205–12 (2014).

28. Ma, W. et al. Fine-scale chromatin interaction maps reveal the cis-regulatory landscape of human lincRNA genes. Nat Methods 12, 71–8 (2015).

29. Hsieh, T.H. et al. Mapping Nucleosome Resolution Chromosome Folding in Yeast by Micro-C. Cell 162, 108–19 (2015).

30. Fortin, J.P. & Hansen, K.D. Reconstructing A/B compartments as revealed by Hi-C using long-range correlations in epigenetic data. Genome Biol 16, 180 (2015).

31. Huang, J., Marco, E., Pinello, L. & Yuan, G.C. Predicting chromatin organization using histone marks. Genome Biol 16, 162 (2015).

32. Chen, Y., Wang, Y., Xuan, Z., Chen, M. & Zhang, M.Q. De novo deciphering three-dimensional chromatin interaction and topological domains by wavelet transformation of epigenetic profiles. Nucleic Acids Res 44, e106 (2016).

33. Zhu, Y. et al. Constructing 3D interaction maps from 1D epigenomes. Nat Commun 7, 10812 (2016).

34. Whalen, S., Truty, R.M. & Pollard, K.S. Enhancer–promoter interactions are encoded by complex genomic signatures on looping chromatin. Nature Genetics 48, 488–96 (2016).

35. He, C., Wang, X. & Zhang, M.Q. Nucleosome eviction and multiple co-factor binding predict estrogen-receptor-alpha-associated long-range interactions. Nucleic Acids Res 42, 6935–44 (2014).

36. Pennings, S., Muyldermans, S., Meersseman, G. & Wyns, L. Formation, stability and core histone positioning of nucleosomes reassembled on bent and other nucleosome-derived DNA. J Mol Biol 207, 183–92 (1989).

37. Kornberg, R.D. & Stryer, L. Statistical distributions of nucleosomes: nonrandom locations by a stochastic mechanism. Nucleic Acids Res 16, 6677–90 (1988).

38. Wang, G.G., Allis, C.D. & Chi, P. Chromatin remodeling and cancer, Part II: ATP-dependent chromatin remodeling. Trends Mol Med 13, 373–80 (2007).

39. Gaffney, D.J. et al. Controls of nucleosome positioning in the human genome. PLoS Genet 8, e1003036 (2012).

40. Tonks, L. The complete equation of state of one two and three-dimensional gases of hard elastic spheres. Phys Rev 50, 955–963 (1936).

41. Kundaje, A. et al. Ubiquitous heterogeneity and asymmetry of the chromatin environment at regulatory elements. Genome Res 22, 1735–47 (2012).

42. Nie, Y., Cheng, X., Chen, J. & Sun, X. Nucleosome organization in the vicinity of transcription factor binding sites in the human genome. BMC Genomics 15, 493 (2014).

43. Consortium, T.E.P. An integrated encyclopedia of DNA elements in the human genome. Nature 488, 57–74 (2013).

44. Davey, C. & Allan, J. Nucleosome positioning signals and potential H-DNA within the DNA sequence of the imprinting control region of the mouse Igf2r gene. Biochim Biophys Acta 1630, 103–16 (2003).

45. Gasiorek, J.J. & Blank, V. Regulation and function of the NFE2 transcription factor in hematopoietic and non-hematopoietic cells. Cell Mol Life Sci 72, 2323–35 (2015).

46. Hubner, M.R. & Spector, D.L. Chromatin dynamics. Annu Rev Biophys 39, 471–89 (2010).

47. Genomes Project, C. et al. A global reference for human genetic variation. Nature 526, 68–74 (2015).

48. Fullwood, M.J. & Ruan, Y. ChIP-based methods for the identification of long-range chromatin interactions. J Cell Biochem 107, 30–9 (2009).

49. He, B., Chen, C., Teng, L. & Tan, K. Global view of enhancer-promoter interactome in human cells. Proc Natl Acad Sci U S A 111, E2191–9 (2014).

50. Chevereau, G., Palmeira, L., Thermes, C., Arneodo, A. & Vaillant, C. Thermodynamics of intragenic nucleosome ordering. Phys Rev Lett 103, 188103 (2009).

51. Landt, S.G. et al. ChIP-seq guidelines and practices of the ENCODE and modENCODE consortia. Genome Res 22, 1813–31 (2012).

52. Thurman, R.E. et al. The accessible chromatin landscape of the human genome. Nature 489, 75–82 (2012).

53. Rozowsky, J. et al. AlleleSeq: analysis of allele-specific expression and binding in a network framework. Mol Syst Biol 7, 522 (2011).

54. Chen, W. et al. Improved nucleosome-positioning algorithm iNPS for accurate nucleosome positioning from sequencing data. Nat Commun 5, 4909 (2014).

55. Zhang, Y., Shin, H., Song, J.S., Lei, Y. & Liu, X.S. Identifying positioned nucleosomes with epigenetic marks in human from ChIP-Seq. BMC Genomics 9, 537 (2008).

56. van Berkum, N.L. et al. Hi-C: a method to study the three-dimensional architecture of genomes. J Vis Exp 6, 1869 (2010).

57. Yaffe, E. & Tanay, A. Probabilistic modeling of Hi-C contact maps eliminates systematic biases to characterize global chromosomal architecture. Nat Genet 43, 1059–65 (2011).

58. Mozziconacci, J. & Koszul, R. Filling the gap: Micro-C accesses the nucleosomal fiber at 100–1000 bp resolution. Genome Biol 16, 169 (2015).

59. Imakaev, M. et al. Iterative correction of Hi-C data reveals hallmarks of chromosome organization. Nat Methods 9, 999–1003 (2012).

60. Williamson, I. et al. Spatial genome organization: contrasting views from chromosome conformation capture and fluorescence in situ hybridization. Genes Dev 28, 2778–91 (2014).

61. Li, Q., Peterson, K.R., Fang, X. & Stamatoyannopoulos, G. Locus control regions. Blood 100, 3077–86 (2002).

62. Dostie, J. et al. Chromosome Conformation Capture Carbon Copy (5C): a massively parallel solution for mapping interactions between genomic elements. Genome Res 16, 1299–309 (2006).

63. Sanyal, A., Lajoie, B.R., Jain, G. & Dekker, J. The long-range interaction landscape of gene promoters. Nature 489, 109–13 (2012).

64. Tolhuis, B., Palstra, R.J., Splinter, E., Grosveld, F. & de Laat, W. Looping and interaction between hypersensitive sites in the active beta-globin locus. Mol Cell 10, 1453–65 (2002).

65. Bailey, S.D. et al. ZNF143 provides sequence specificity to secure chromatin interactions at gene promoters. Nat Commun 2, 6186 (2015).

66. Fu, Y., Sinha, M., Peterson, C.L. & Weng, Z. The insulator binding protein CTCF positions 20 nucleosomes around its binding sites across the human genome. PLoS Genet 4, e1000138 (2008).

